# Metabolic profiling of licensed mesenchymal stromal cells reveals a disconnect between glycolytic fitness and immunomodulatory responsiveness

**DOI:** 10.64898/2026.09.27.754820

**Authors:** Haider Bilal, Brendon T. DeGroot, Lorena R. Braid

## Abstract

Mesenchymal stromal cells (MSCs) are promising cell therapeutics, but their development as off-the-shelf products requires cryopreservation, which impairs cell function after thaw. Inflammatory licensing enhances MSC therapeutic attributes, yet protocols vary considerably and little is known about how these variables affect MSC metabolism or post-thaw recovery. We characterized mitochondrial and glycolytic flux in umbilical cord (UC) and bone marrow (BM) MSCs licensed with interferon-γ (IFN-γ), tumor necrosis factor-α (TNF-α), or both. Brief exposure to either cytokine shifted MSC metabolism toward glycolysis and reduced mitochondrial respiration, whereas prolonged exposure led to heterogeneous responses. Conversely, combined IFN-γ and TNF-α increased glycolytic flux and respiratory capacity in most donors, with responses detectable after 2 hours and at concentrations below those conventionally used. Cryopreservation impaired aerobic metabolism, followed by donor-dependent recovery. Preconditioning with combined IFN-γ and TNF-α partially improved post-thaw metabolism, particularly in BM MSCs, and enhanced early responsiveness to inflammatory challenge, particularly in UC MSCs, as detected by *IDO* and candidate biomarkers *CTSS* and *C1S*. However, donors with improved post-thaw metabolism did not consistently show increased biomarker responses. These findings demonstrate that metabolic fitness and inducible biomarkers capture complementary aspects of post-thaw MSC quality and may inform preconditioning strategies for cryopreserved MSC products.

## INTRODUCTION

Mesenchymal stromal cells (MSCs) are promising cell therapy candidates due to their immunomodulatory, regenerative, and paracrine properties which confer reparative benefits for a spectrum of clinical applications^1–9^. MSCs doses are prepared from active cell cultures (fresh) or thawed from cryopreserved stocks ^3,7,10^. While stockpiled MSCs are more feasible for commercialization, numerous studies have shown that their function is compromised after thaw ^2,7,11–14^. Stress factors such as plasma membrane alterations, cytoprotectant cytotoxicity, heat shock, and metabolic uncoupling are freeze-thaw induced stressors and that impact the viability, cellular attachment, immunomodulation, and metabolism of MSCs immediately after thaw^6,7,15,16^.

Cryopreservation is a critical step in the MSC pipeline to facilitate stockpiles of qualified MSC products for allogeneic, on-demand applications, and to retain master stocks of limited patient-specific MSCs ^3,4,10–12,17^. As such, there is a critical need to better understand the functional impairments of MSCs immediately after thaw and to identify strategies to mitigate these deficits and/or to accelerate recovery after thaw. Studies have shown that thawed MSCs can be “refreshed” by returning them to cell culture before administration, but this process abrogates the economic and practical benefits of “off-the-shelf” MSC therapies. However, the culture time required to functionally recover MSCs is not yet clearly defined, and some groups even report that thawed MSCs perform just as well as their fresh counterparts^18–21^. This lack of consensus has culminated in the use of thawed MSCs in clinical trials with significant variability in the time from thaw to administration^2,3,11,13,17^, making it difficult to use retrospective analysis to determine best practices in recovery time and processes for cryopreserved MSC products. Several strategies are being explored to improve MSC function, including cultivating the cells under hypoxia, licensing with inflammatory cytokines, specialized cryopreservation media, and embedding the cells in biomaterials like hydrogels^22–26^. Preconditioning MSCs with cytokines like IFN-γ and TNFα before cryopreservation has been shown to improve their function after thaw by “priming” the MSCs to respond to the inflammatory milieu in the recipient^9,11,27–29^. In addition, IFN-γ and TNF-α stimulation are known to direct MSCs into glycolysis through PI3K-AKT signalling^30^. In glycolysis, MSCs upregulate expression of the immunomodulatory factors IDO and TSG-6, thus enhancing their anti-inflammatory benefits^31^. Glycolysis has also been shown to be important for MSC survival after implantation, as well as enhanced therapeutic efficacy and immunomodulation^32,33^.

In general, metabolism is thought to be an important driver of MSC function, both from its production of metabolites and energy, and its role in cell signalling. For example, the well-known readout of MSC immunomodulation and T-cell suppression: via indoleamine 2,3-dioxygenase (IDO), is an enzyme involved in tryptophan-kynurenine metabolism^34^. However, there is conflicting evidence about how cytokine stimulation influences MSC metabolism. For example, Liu et al reported that IFN-γ licensing promoted a shift to glycolysis in bone marrow (BM) MSCs by increasing their extracellular acidification rate (ECAR) and expression of glycolytic genes without affecting their oxygen consumption rate (OCR)^32^. Furthermore, rotenone-induced suppression of OXPHOS prior to IFN-γ stimulation was shown to augment IDO expression and activity beyond that observed with stimulation alone, implying an inverse correlation between OXPHOS and IDO-dependent immunomodulation^32^. By contrast, Yao et al. observed that IFN-γ-activated umbilical cord (UC) MSCs increased OCR, decreased ECAR and had no effect on glucose uptake^35^. In this study, rotenone pre-treatment reduced IDO and PD-L1 protein levels, suggesting that OXPHOS promotes UC MSC immunomodulation^35^. Licensing MSCs with a cocktail of IFN-γ and TNF-α has been reported to increase glycolytic flux^26,29^ and conversely to decrease metabolic flux^30^.

These disparate findings suggest that metabolic responses to cytokines like IFN-γ and TNF α may be influenced by intrinsic and extrinsic factors, including MSC tissue of origin, donor, culture media, atmospheric conditions, and cytokine identity, dose, and duration of licensing. Consistent with this hypothesis, it has already been shown that MSCs cultivated in media supplemented with human platelet lysate (hPL) have higher metabolic activity compared to MSCs cultured in FBS-supplemented media^36^. Most studies culture MSCs in 5% CO_2_ and ambient oxygen tension (∼20%), although oxygen saturation in tissues (physioxia) is typically 3-5%. Among other differences, MSCs cultured in < 5% O_2_ exhibit higher levels of anaerobic glycolysis than MSCs in ambient oxygen^37,38^. Indeed, the 4-fold higher level of oxygen in ambient conditions may bias MSCs towards OXPHOS, and data from these experiments may not accurately reflect the metabolic flux of preconditioned-MSCs when infused into tissues^39^. In addition, cytokine licensing is performed using a wide range of protocols. Published reports have inoculated MSCs with 5 to 40 ng/ml of IFN-γ and/or TNF-α for 16 to 72 hours^11,28,31,33,35,40,41^. It has already been shown that cytokines can have a time and dose dependent effect on MSCs^42–46^. For example, TNFα reportedly promotes osteogenic differentiation of BM MSCs at lower doses, but inhibits osteogenesis and promotes apoptosis at higher concentrations^43,44^. Although prior studies have examined how IFN-γ dose and stimulation duration affect IDO expression and function^46,47^, no comprehensive analysis across multiple MSC donors has yet evaluated metabolic flux and immunomodulatory gene expression under single or dual IFN-γ + TNF-α stimulation.

The aim of this study is to reconcile the reported disparities and knowledge gaps by completing side-by-side metabolic analyses of 3 donor populations each of fresh and cryopreserved UC and BM MSCs. We characterize the resting metabolic state of each MSC lot, assess the time to metabolic recovery after thaw, and the metabolic impacts of licensing with IFN-γ and TNF-α at doses ranging from 0.1 to 10 ng/ml and for durations of 2-24 hours. We also study the benefits of cytokine preconditioning before cryopreservation with respect to metabolic recovery and ability to respond to inflammatory challenge. Together, these data reveal new insights into the intrinsic metabolic capacity of individual donor MSC populations and challenge the current accepted paradigm that metabolic status correlates with functional potency.

## MATERIALS AND METHODS

### Cell culture

UC MSCs and BM MSCs (Tissue Regeneration Therapeutics (TRT) Inc., CAN, and RoosterBio Inc, USA) were thawed at passage 2 (P2) according to TRT’s proprietary standard operating procedures and expanded in replicate using xeno-free human platelet lysate (hPL)-supplemented media (XF-hPL; RoosterBio Inc, USA) at 37°C, 5% CO_2_, and 5% O_2_ in T75 flasks (Greiner Bio-One) until 80% confluence. Flasks were pre-coated with human placental Collagen IV (Sigma Aldrich #C5533) diluted in Dulbecco’s phosphate-buffered saline without calcium or magnesium (DPBS^-/-^; Sigma-Aldrich, St. Louis, MO, USA) at a density of 2.67 ug/cm^2^ for 2 hours at room temperature. Once confluent, cells of one replicate flask were activated upon a 2-hour exposure to human IFN-γ (10 ng/mL) and/or TNF-α (10 ng/mL) (PeproTech, Rocky Hill, NJ, USA) under the incubation conditions mentioned above. Cells were then washed with DPBS^-/-^ before being enzymatically detached by TrypLE Select (ThermoFisher Scientific, Waltham, MA, USA). Once dissociated, an equal volume of media was added to the cell suspension and cells were counted with a Millipore Scepter cell counter using 60 mm probes (Millipore, Billerica, MA, USA). Cells were then centrifuged at 149*g* for 5 minutes and the cell pellet resuspended in 1 mL of fresh media per aliquot.

### Cryopreservation and thawing of MSCs

To examine whether activation of MSCs with IFN-γ + TNF-α before freezing enhanced metabolic recovery, parallel cultures of stimulated and unstimulated MSCs were frozen at P3 at a concentration of 1 to 2×10^6^ cells/mL using equal volumes of EZ-CPZ (INCELL, San Antonio, TX, USA) and media. Cells were placed in a freezing container (CoolCell® FTS30; Corning®, NY, USA) with a cooling rate of -1 °C/minute until -80°C internal temperature was reached. MSC aliquots were transferred into liquid nitrogen for at least 24 hours to achieve a state of cryopreservation. A few days prior to experimentation, age and donor-matched MSCs (P2) were thawed and expanded to serve as a cultured control group. On the day of experimentation, cryopreserved MSCs were thawed using the protocol described above. Once thawed, cells were centrifuged at 149*g* for 5 minutes and the cell pellet resuspended in culture media. Age-matched fresh MSCs (P3) were harvested alongside thawed cells, centrifuged, and resuspended in complete media for metabolic analysis.

### Metabolic analysis

MSC energetics were assessed by measuring oxygen consumption rate (OCR), extracellular acidification rate (ECAR), and NAD(P)H-dependent oxidoreductase activity. OCR and ECAR were measured using the Mito Stress Test on the Seahorse XF Pro/e96 analyzer (Agilent, Santa Clara, CA, USA). To evaluate the metabolic effects of IFN-γ + TNF-α (PeproTech, Rocky Hill, NJ, USA), fresh MSCs were seeded at 12,500 cells/well in Collagen IV-coated Seahorse XF96 V3 PS plates (part #101) 24 hours before cytokine stimulation. Cells were then dosed with IFN-γ and/or TNF-α at various concentrations (0.1–10 ng/ml) for 2, 4, 8, or 24 hours. In parallel, fresh and thawed age-matched UC and BM MSCs were seeded at 6,250, 12,500 or 25,000 cells per well for the 6-, 24-, or 48-hour post-thaw or suspension groups, respectively. After cytokine stimulation or time post-thaw was complete, the Mito Stress Test was conducted as per the manufacturer’s instructions.

The sensor cartridge was hydrated overnight in XF Calibrant (Cat. No. 100840-000) at 37 °C in a non-CO₂ incubator. On the day of the assay, culture media was replaced with Seahorse XF DMEM Assay Medium, pH 7.4 (Cat. No. 103575-100), supplemented with 10 mM glucose (Cat. No. 103577-100), 1 mM sodium pyruvate (Cat. No. 103578-100), and 2 mM L-glutamine (Cat. No. 103579-100). The wells were rinsed 3 times with the Seahorse medium to ensure full elimination of the hPL-supplemented culture medium.

Once the test was complete, cells were washed 3 times with 100 µL/well of DPBS for 5 minutes, then fixed in 100 µLof 4% paraformaldehyde diluted in DPBS for 10 minutes with gentle rocking. Nuclei were stained with Hoechst 33342 (ThermoFisher Scientific) at a 1:1000 dilution in DPBS for 10 minutes at room temperature. Cells were again washed thrice with DPBS. The entire well was imaged using a Nikon Eclipse Ti2 high-content fluorescence microscope. Images were acquired with Nikon JOBS software using continuous autofocus and automatically saved as .nd2 files. The NIS software was used quantify the number of nuclei per well identified from the 365 nm channel. All data from the Seahorse Analyzer was then normalized to the number of nuclei per well.

NAD(P)H-dependent oxidoreductase enzyme activity was measured using CCK-8 assay kits (Sigma-Aldrich) at 0-, 2-, 4-, 8-, 24-, and 48-hours post-suspension or thaw according to manufacturer’s instructions. This assay quantifies the NAD(P)H-dependent reduction of the water-soluble tetrazolium salt (WST-8) to a water-soluble formazan dye. The absorbance of formazan is proportional to the concentration of NAD(P)H, an important cofactor in enzymatic reactions essential for cellular metabolism and mitochondrial function. For each condition, MSCs were seeded in a 96-well plate at a density of 10,000 cells/well in 100 mL of media with four replicate wells for evaluation at each time point. Once seeded, cells were incubated at 37°C, 5% CO_2_ for the various durations listed above. At each time point, 10 mL of CCK-8 reagent was added to each well of the plate and incubated at 37°C, 5% CO_2._ After a 2-hour incubation period, absorbance was measured at 450 nm using a BioTek Synergy HTX microplate reader (Agilent).

### RT-qPCR and RNA Extraction

RT-qPCR was performed using the Applied Biosystems Power SYBR green RNA to CT 1-step kit with 50 ng total RNA. Data was normalized to the mean expression of ACTB, GAPDH and YWHAZ reference genes. RNA was extracted using the Norgen Total RNA Purification kit, from freeze-thawed MSC cell lysate stored in RNA-Later (Norgen). The CFX96 RT-qPCR machine (Bio-Rad Laboratories) was used to run the assay, and normalized gene expression was calculated using the native CFX96 software (CFX Maestro 2.3)

### Statistical analysis

Data was analyzed using GraphPad Prism 9.0 software. An unpaired two-sided t test was used to determine significance between the means of two groups, and a one-way ANOVA with multiple comparisons was used to compare multiple groups simultaneously. A p value < 0.05 was deemed statistically significant.

## RESULTS

### Pro-inflammatory cytokine licensing transiently shifts MSC metabolism to glycolysis while suppressing oxidative metabolism

To assess the metabolic consequences of inflammatory licensing, BM and UC MSCs from three donors each were treated with 1 or 10 ng/ml IFN-γ or TNF-α for 2 or 24 h. Mitochondrial function and metabolic flux were assessed using the Seahorse XF Cell Mito Stress Test, including basal and maximal respiration, spare respiratory capacity, ATP-linked respiration, extracellular acidification rate (ECAR), and the OCR/ECAR ratio (Fig. 1). Because metabolic responses varied among donor populations, each donor was analyzed independently rather than pooling data.

**Figure 1.**
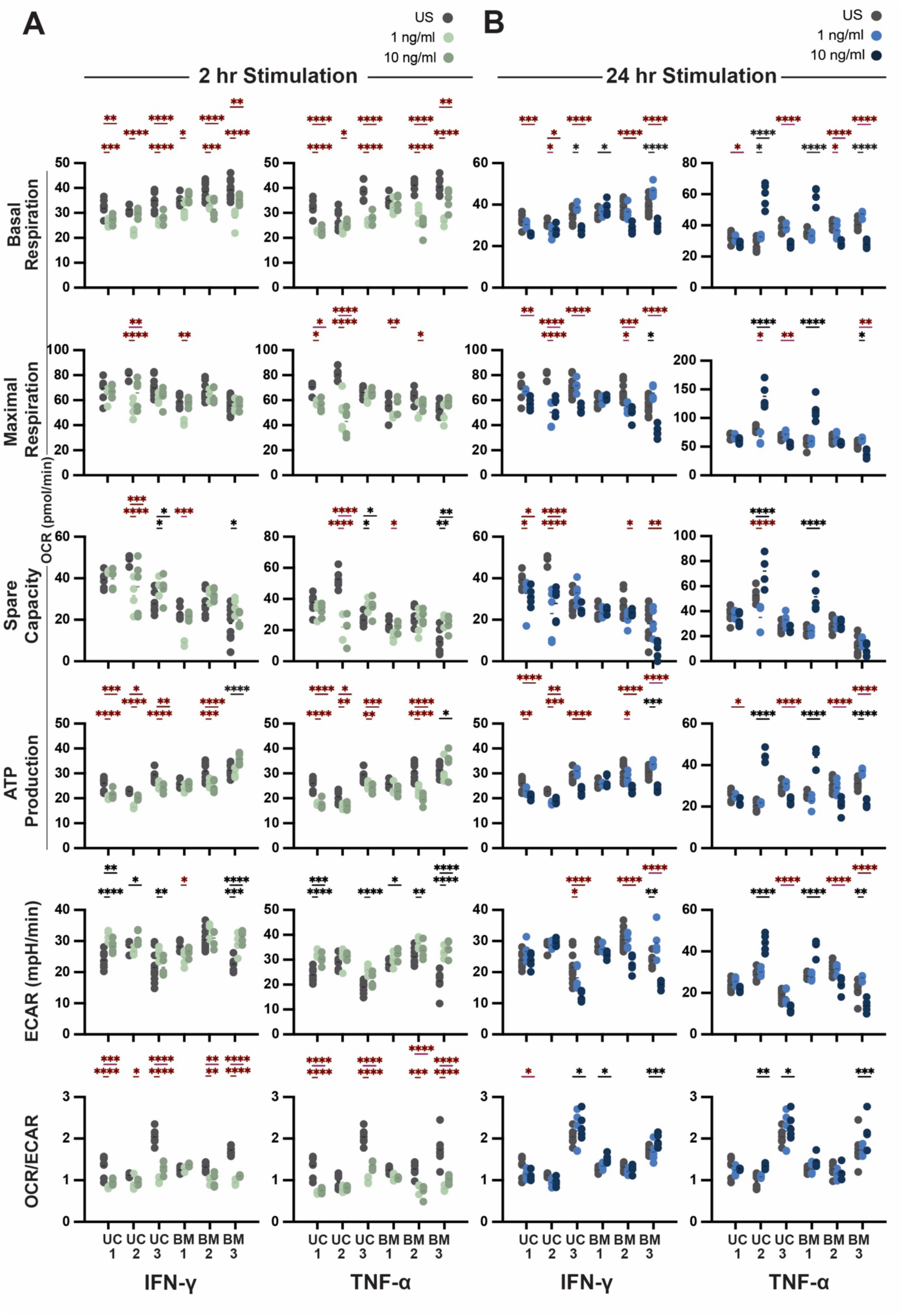
Dose and duration of IFN-γ or TNF-α licensing affect the metabolic status of UC and BM MSCs. Metabolic flux was measured in three umbilical cord (UC1–UC3) and three bone marrow (BM1–BM3) MSC donor populations using the Seahorse XF Mito Stress Test following licensing with 1 or 10 ng/ml IFN-γ or TNF-α for (A) 2 h or (B) 24 h. (A) After 2 h, glycolytic flux increased in most donors, whereas basal respiration, maximal and spare respiratory capacity, and ATP-linked respiration generally decreased, although responses varied by donor and cytokine. Accordingly, the OXPHOS/glycolysis ratio decreased in 4/6 donors following either cytokine. (B) At 24h, metabolic responses were more variable, with fewer donors showing increased glycolytic flux and less consistent changes in respiratory parameters, indicating that the glycolytic shift observed at 2h diminished with prolonged licensing. Grey dots indicate unstimulated (US) MSCs; light and dark green (A) or blue (B) dots indicate 1 and 10 ng/ml cytokine, respectively. Dark red asterisks indicate statistically significant decreases and black asterisks indicate statistically significant increases.

Short-term cytokine licensing generally shifted MSC metabolism toward increased glycolysis and reduced oxidative phosphorylation (Fig. 1A). After 2 h, TNF-α increased ECAR in 5/6 donors, with UC2 remaining unchanged, whereas IFN-γ increased ECAR in 4/6 donors, with BM1 and BM2 remaining unchanged or decreasing. This glycolytic response was less apparent after 24 h (Fig. 1B). TNF-α increased ECAR in only a subset of donors, with responses varying by dose, while IFN-γ increased ECAR only in BM3 at the lower dose.

Changes in mitochondrial respiration were similarly dependent on cytokine dose and exposure time. After 2 h, low-dose IFN-γ decreased basal respiration in all six donors, whereas the high dose decreased basal respiration in 4/6 donors (Fig. 1A). TNF-α also decreased basal respiration in most donors at 2 h. After 24 h, responses became more heterogeneous: IFN-γ decreased basal respiration in UC2 and BM3 at both doses and in UC1, UC3, and BM2 at the higher dose, but increased basal respiration in UC3 and BM3 at the lower dose and BM1 at the higher dose (Fig. 1B). TNF-α produced similarly variable responses after 24 h, including decreased basal respiration in BM2 at both doses and UC3 and BM3 at the higher dose, but increased respiration in UC2, BM1, or BM3 under other dose conditions.

Maximal respiration and spare respiratory capacity also declined in a subset of donors. After 2 h of IFN-γ licensing, both parameters decreased in UC2 and BM1, whereas TNF-α decreased them in UC1, UC2, BM1, and BM2 (Fig. 1A). After 24 h of IFN-γ licensing, maximal and spare respiratory capacity were reduced in 4/6 donors, while responses to TNF-α were more heterogeneous (Fig. 1B). ATP-linked respiration was reduced in all three UC donors and BM2 after 2 h of either cytokine. At 24 h, IFN-γ reduced ATP-linked respiration in all UC donors, whereas responses to TNF-α varied among donors and doses.

Consistent with these changes, 2 h of cytokine licensing generally decreased the OCR/ECAR ratio, reflecting a shift toward glycolytic metabolism (Fig. 1A). This effect was diminished after 24 h, when metabolic responses were more heterogeneous (Fig. 1B). Collectively, these data demonstrate that IFN-γ and TNF-α alter MSC metabolic state, with the magnitude and direction of the response dependent on exposure duration, cytokine dose, and donor population.

### Combined IFN-γ and TNF-α boosts MSC metabolism in a time- and dose-dependent manner

To determine the metabolic effects of combined cytokine licensing, the same six MSC donor populations were stimulated with IFN-γ and TNF-α while varying either the duration or concentration of stimulation (Fig. 2). At 10 ng/ml, dual stimulation increased maximal respiration and spare respiratory capacity in most donors, although the magnitude and timing of the response varied among donors (Fig. 2A). ECAR increased after 2 h in 5/6 donors, with UC3 showing a more time-dependent response (Fig. 2A). In contrast, ATP-linked respiration decreased in UC1, UC2, and BM3 following dual stimulation, whereas BM1 increased (Fig. 2A). Consistent with increased glycolytic flux, the OCR/ECAR ratio generally decreased following stimulation, although the timing of this response varied among donors.

**Figure 2.**
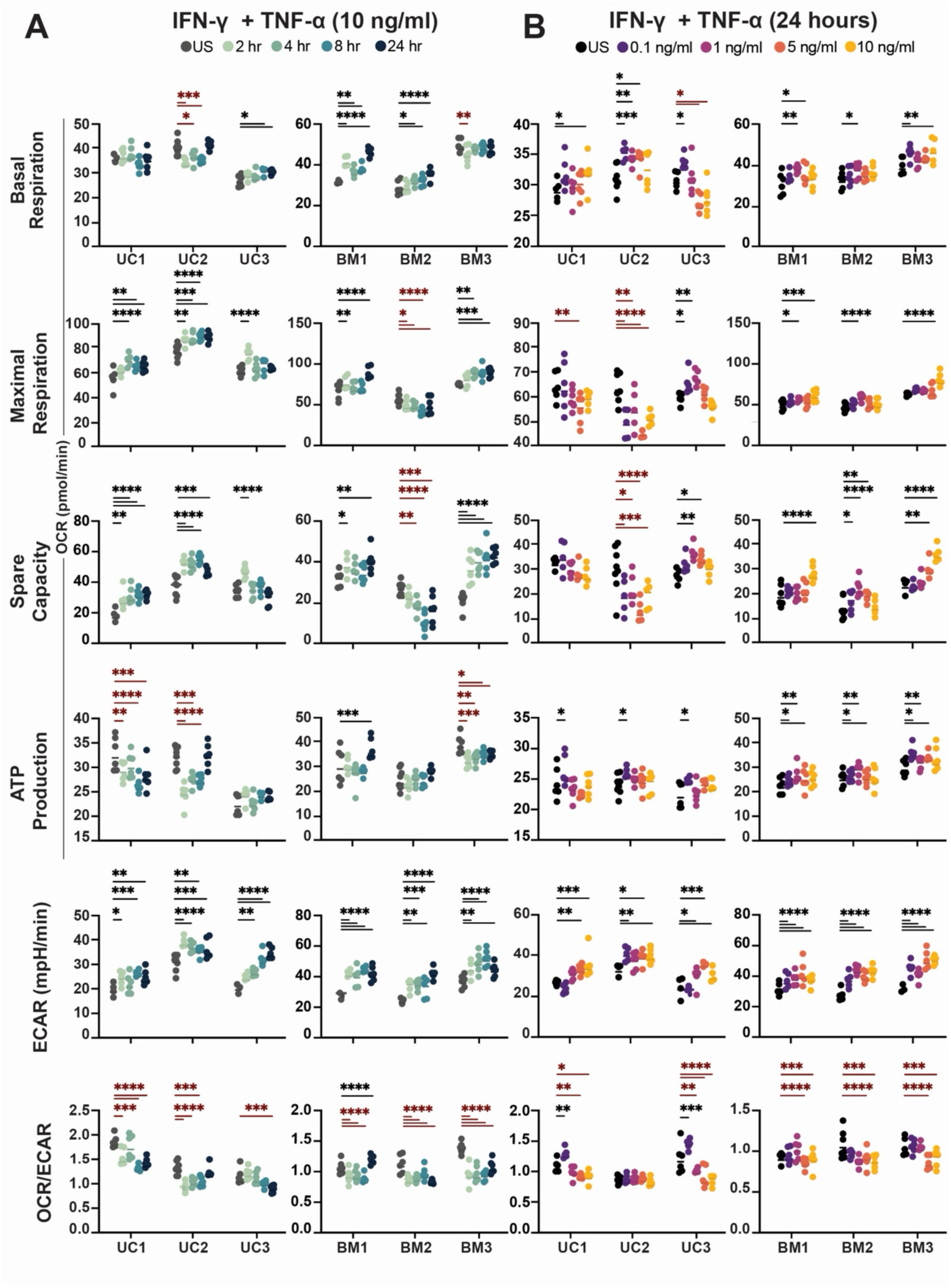
Duration and dose of combined IFN-γ and TNF-α stimulation alter aerobic and glycolytic metabolism in UC and BM MSCs. Metabolic flux was measured in three umbilical cord (UC1–UC3) and three bone marrow (BM1–BM3) MSC donor populations using the Seahorse XF Mito Stress Test. (A) MSCs were stimulated with 10 ng/ml IFN-γ + TNF-α for 2, 4, 8, or 24 h. ECAR generally increased following dual stimulation, while maximal respiration and spare respiratory capacity increased in most donors, with responses varying by donor and duration. ATP-linked respiration showed more variable responses. (B) MSCs were stimulated for 24 h with 0.1, 1, 5, or 10 ng/ml IFN-γ + TNF-α. Metabolic responses were dose- and donor-dependent, with changes in basal and maximal respiration, spare respiratory capacity, ATP-linked respiration, and ECAR across the concentration range. The OCR/ECAR ratio generally decreased with dual stimulation, consistent with a relative shift toward glycolytic metabolism. In (A), grey indicates unstimulated (US), and progressively darker green/blue dots indicate 2, 4, 8, and 24 h stimulation. In (B), black indicates US, and purple, magenta, orange, and yellow indicate 0.1, 1, 5, and 10 ng/ml, respectively. Dark red asterisks indicate statistically significant decreases and black asterisks indicate statistically significant increases.

Metabolic responses after 24 h of dual stimulation were also concentration- and donor-dependent (Fig. 2B). In UC MSCs, low-dose stimulation increased basal respiration and ATP-linked respiration, whereas higher concentrations produced variable effects on basal respiration (Fig. 2B). ECAR also increased across the UC donors, although the concentration associated with this response differed among donors (Fig. 2B). In BM MSCs, ECAR increased at low cytokine concentrations, while increases in basal respiration were observed at higher concentrations in BM1 and BM2 and at lower concentrations in BM3 (Fig.2B). Maximal respiration and spare respiratory capacity generally increased with cytokine concentration in BM1 and BM3, whereas BM2 showed a different dose-response pattern (Fig. 2B). Overall, combined IFN-γ and TNF-α licensing altered both oxidative and glycolytic metabolism, with responses dependent on cytokine concentration, stimulation duration, and donor population (Fig. 2).

### Thawed MSCs exhibit impaired metabolism that is partially mitigated by cytokine preconditioning, particularly in BM MSCs

Based on the metabolic response to combined cytokine licensing, MSCs were preconditioned with 10 ng/ml each of IFN-γ and TNF-α for 2 h before cryopreservation. This condition was selected as the shortest tested duration that increased metabolic flux across BM and UC MSCs (Fig. 2A). Following thaw, metabolic recovery was assessed after 6, 24, and 48 h in culture and compared with fresh MSCs (Fig. 3).

**Figure 3.**
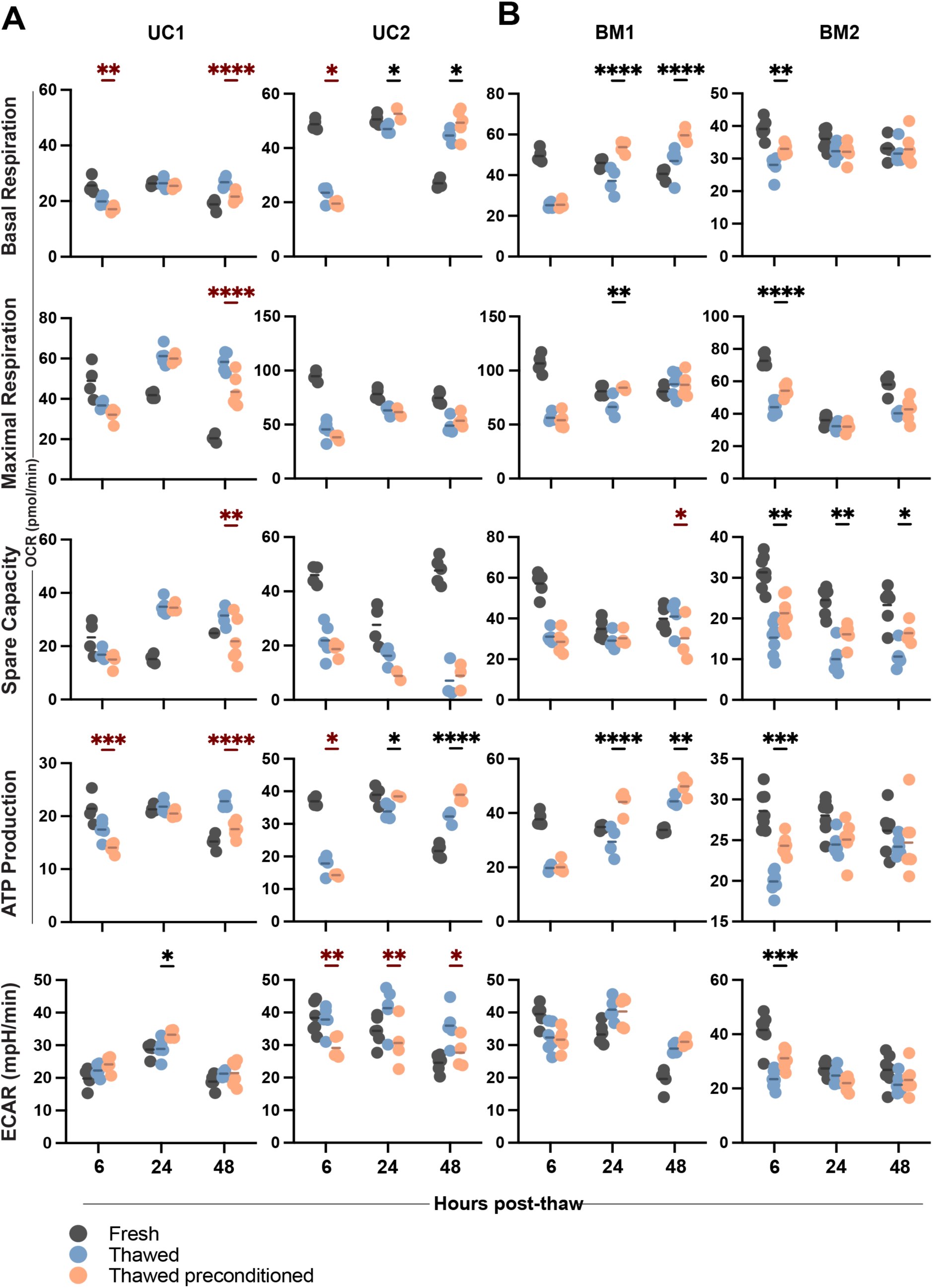
Cytokine preconditioning partially restores aerobic metabolism in thawed MSCs, particularly in BM-derived cells. Metabolic flux was measured in two umbilical cord (UC1–UC2) and two bone marrow (BM1–BM2) MSC donor populations using the Seahorse XF Mito Stress Test. Fresh MSCs and MSCs thawed with or without cytokine preconditioning were recovered in culture for 6, 24, or 48 h before metabolic analysis. (A) In UC MSCs, thawing reduced measures of aerobic metabolism in a donor- and recovery time-dependent manner, with limited improvement following preconditioning. (B) In BM MSCs, preconditioning improved several measures of aerobic metabolism following thaw, although responses varied by donor and recovery time. Grey dots indicate fresh MSCs, blue dots indicate thawed MSCs, and orange dots indicate preconditioned thawed MSCs. Dark red asterisks indicate statistically significant decreases and black asterisks indicate statistically significant increases.

Thawing initially impaired aerobic metabolism in both UC donors (Fig. 3A). At 6 h, basal and maximal respiration, spare respiratory capacity, and ATP-linked respiration were lower in thawed cells than in fresh cells, regardless of preconditioning, whereas ECAR was less affected. Recovery was donor-dependent. In UC1, respiratory parameters recovered substantially by 24 h and were maintained or increased at 48 h. In contrast, UC2 showed persistent impairment in maximal respiration and spare respiratory capacity through 48 h despite recovery of basal respiration. Overall, preconditioning provided limited benefit in UC MSCs, although improvements in selected respiratory parameters were observed in UC2.

BM MSCs also exhibited marked impairment in aerobic metabolism following thaw but showed a more apparent response to preconditioning (Fig. 3B). At 6 h, respiratory parameters were reduced in both BM donors, accompanied by lower ECAR. In BM2, preconditioning improved several measures of aerobic metabolism relative to unconditioned thawed cells at this early time point, whereas little benefit was apparent in BM1. By 24 h, preconditioning improved recovery of respiratory metabolism in BM1, while BM2 showed partial recovery in both thawed groups. At 48 h, most metabolic parameters in BM1 had recovered, whereas BM2 retained deficits in maximal respiration and spare respiratory capacity. Collectively, metabolic impairment following cryopreservation was donor- and time-dependent, with cytokine preconditioning providing a more apparent benefit in BM than UC MSCs.

We next assessed cellular dehydrogenase activity using the tetrazolium-based CCK-8 assay, which measures cellular reducing activity through reduction of WST-8 to a water-soluble formazan product. CCK-8 activity was measured immediately and at 8 and 24 h after thaw in UC2, BM1, and BM2, as well as four additional donors (UC4, UC5, UC6, and BM4) (Fig. S1). Responses varied considerably among donors and recovery time points, with both increases and decreases observed in thawed and preconditioned thawed cells relative to fresh cells. Importantly, preconditioning did not produce a consistent improvement in CCK-8 activity compared with unconditioned thawed cells. Thus, CCK-8 did not consistently discriminate the post-thaw metabolic differences identified by Seahorse metabolic flux analysis.

**Figure S1.**
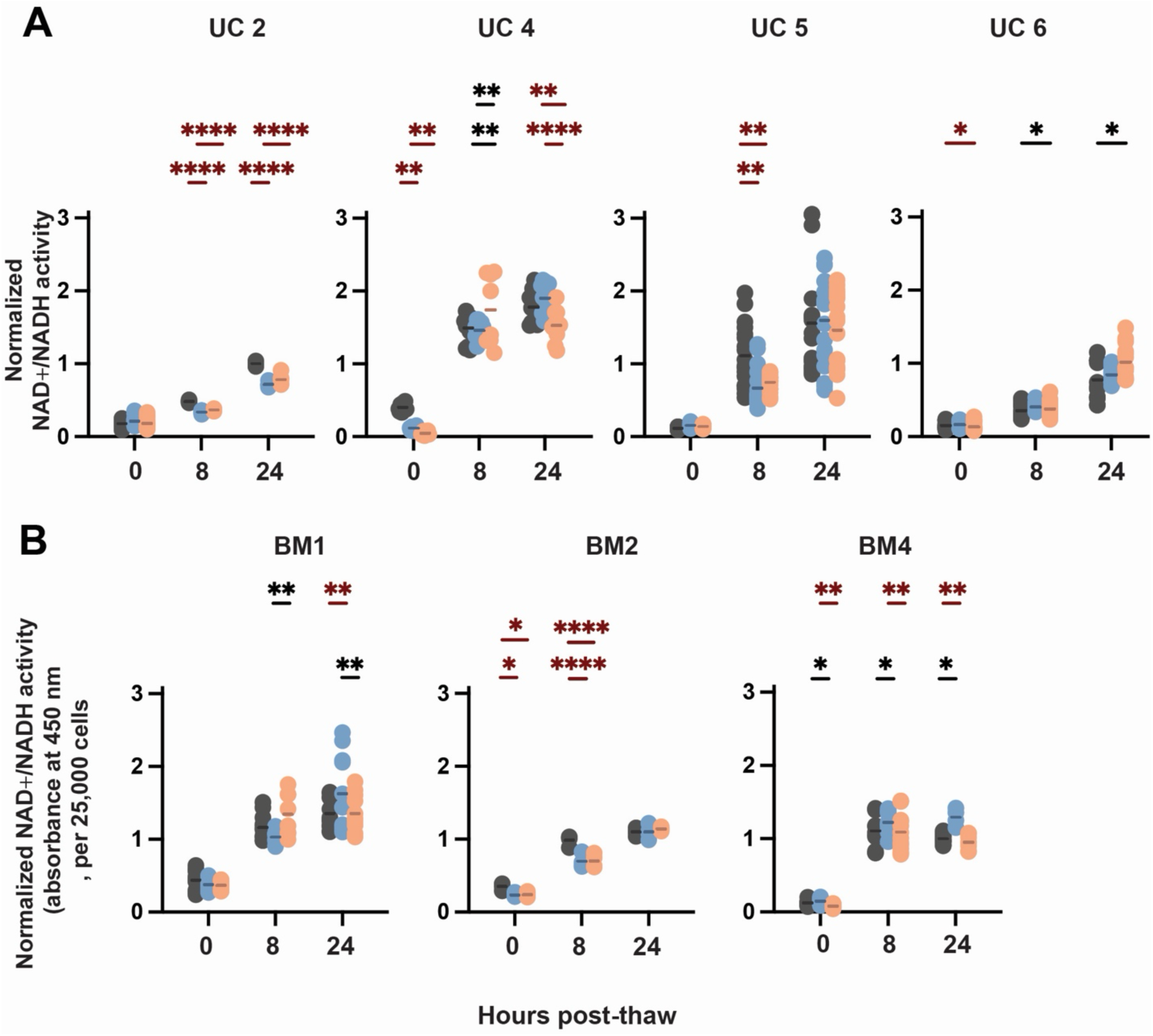
Cytokine preconditioning has variable effects on CCK-8 activity following MSC cryopreservation. CCK-8 activity was measured in four umbilical cord (UC2, UC4–UC6) and three bone marrow (BM1, BM2, BM4) MSC donor populations immediately (0 h), 8 h, and 24 h after thaw. Fresh MSCs were compared with MSCs thawed with or without cytokine preconditioning, and absorbance was normalized to 25,000 cells. Responses varied among donors and recovery times, with no consistent effect of preconditioning across UC or BM MSCs. Grey dots indicate fresh MSCs, blue dots indicate thawed MSCs, and orange dots indicate preconditioned thawed MSCs. Statistical comparisons above each plot are shown for fresh versus thawed, fresh versus preconditioned thawed, and thawed versus preconditioned thawed MSCs. Dark red asterisks indicate statistically significant decreases and black asterisks indicate statistically significant increases.

### Preconditioning enhances expression of established and candidate MSC potency biomarkers

Having observed that preconditioning improved selected parameters of post-thaw metabolism, we next assessed whether preconditioning also enhanced expression of MSC potency-associated genes following inflammatory challenge. Thawed MSCs, with or without preconditioning, were recovered in culture for 6 or 24 h and subsequently challenged with 10 ng/ml IFN-γ + TNF-α for an additional 6 or 24 h. Expression of the established MSC potency marker *IDO*, together with the candidate potency biomarkers *CTSS* and *C1s* previously identified by our group (manuscript in preparation) was then measured by RT-qPCR. Notably, a recent independent study also identified increased *C1s* expression in IFN-γ activated BM MSCs^48^. *CTSS* and *C1s* were included to assess whether their expression captured post-thaw responses to inflammatory challenge alongside the established *IDO* response.

Preconditioning enhanced the response of UC MSCs to inflammatory challenge, with the clearest effects after shorter post-thaw recovery (Fig. 4A, B). Following 6 h of recovery and a 6-h inflammatory challenge, preconditioned UC1 and UC2 MSCs showed higher expression of the established potency marker *IDO* than unconditioned thawed cells (Fig. 4A). Importantly, both candidate biomarkers also detected this response, with *CTSS* and *C1s* similarly increased by preconditioning in both UC donors (Fig. 4A). After 24 h of inflammatory challenge, the benefit of preconditioning remained most evident following the shorter 6-h recovery: *IDO* and *CTSS* remained higher in preconditioned UC2, while *C1s* was higher in preconditioned UC1 and UC2 (Fig. 4B). After 24 h of post-thaw recovery, differences between preconditioned and unconditioned cells were less consistent (Fig. 4). Thus, preconditioning enhanced early post-thaw responsiveness to inflammatory challenge in UC MSCs, and both *CTSS* and *C1s* captured these preconditioning-associated responses alongside *IDO*.

**Figure 4.**
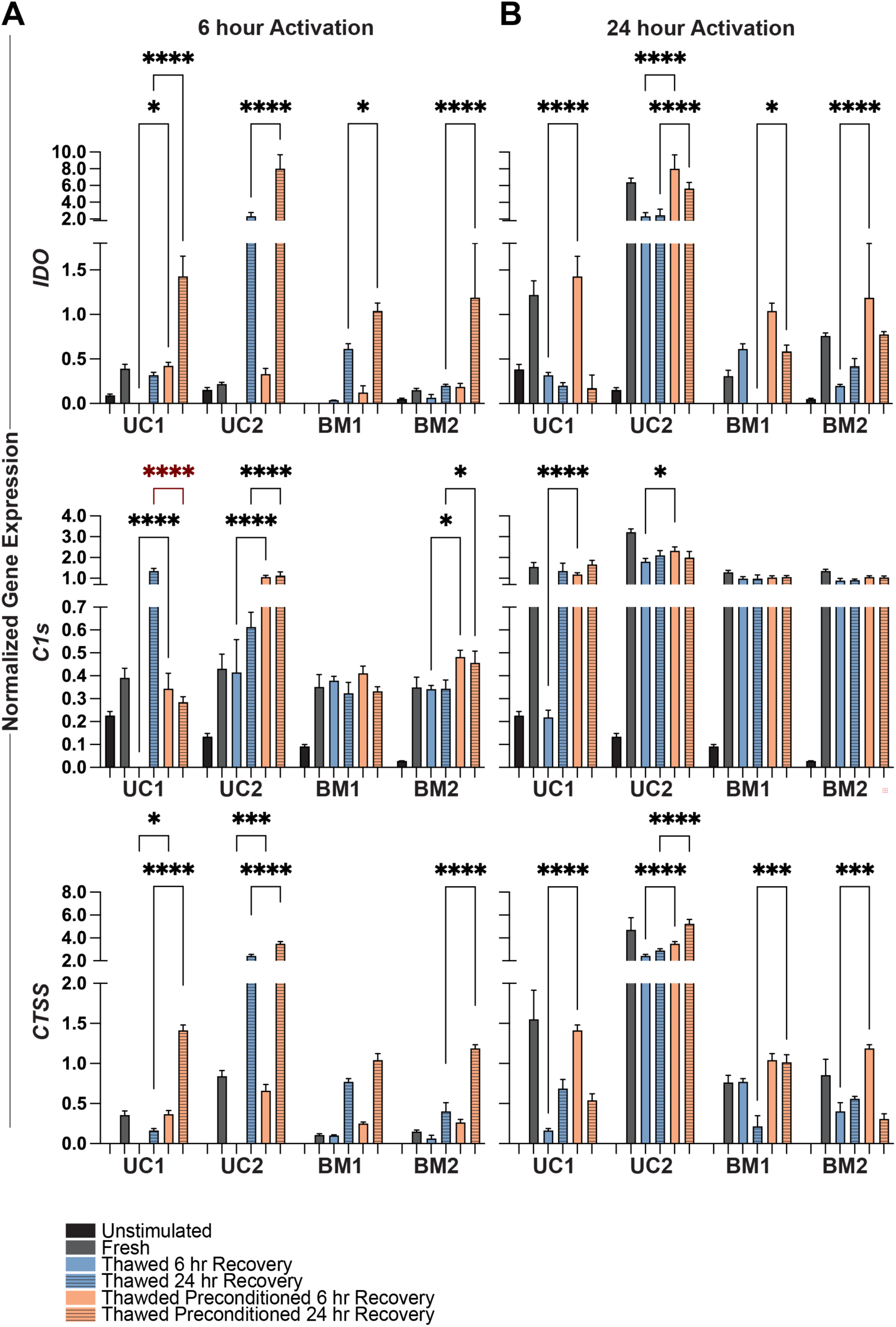
Cytokine preconditioning enhances MSC potency biomarker expression following inflammatory challenge. UC and BM MSCs were thawed and recovered in culture for 6 or 24 h before inflammatory challenge with 10 ng/ml IFN-γ and TNF-α for an additional (A) 6 h or (B) 24 h. Fresh MSCs were challenged under the same conditions. Expression of the potency-associated genes *IDO*, *C1s*, and *CTSS* was quantified by RT-qPCR. Preconditioning enhanced biomarker expression under selected conditions, with responses varying by donor, gene, recovery time, and duration of inflammatory challenge. Effects were generally more consistent in UC than BM MSCs and were detectable as early as 6 h after thaw. Bars indicate unstimulated (black), fresh stimulated (grey), thawed after 6 h (blue) or 24 h (striped blue) recovery, and preconditioned thawed after 6 h (orange) or 24 h (striped orange) recovery. Dark red asterisks indicate statistically significant decreases and black asterisks indicate statistically significant increases.

Preconditioning also enhanced post-thaw inflammatory responses in BM MSCs, but the effects were more dependent on donor, recovery time, and gene (Fig. 4A, B). Following a 6-h inflammatory challenge, preconditioning increased *IDO* under selected conditions (Fig. 4A), with the clearest response in BM2 after 24 h of recovery (Fig. 4B). *CTSS* also detected a preconditioning-associated response in BM2, whereas *C1s* showed little difference between preconditioned and unconditioned thawed BM cells (Fig. 4A, B). Following a 24-h challenge, preconditioning increased *IDO* in BM1 after 24 h of recovery and in BM2 after 6 h of recovery, while *CTSS* responses differed between donors, occurring most clearly after 24 h of recovery in BM1 and 6 h in BM2 (Fig. 4A, B). *C1s* remained similar between preconditioned and unconditioned cells under most BM conditions (Fig. 4A, B). Thus, preconditioning enhanced selected post-thaw responses in BM MSCs, but unlike in UC MSCs, *CTSS* and particularly *C1s* did not consistently report these changes across BM donors and conditions.

Overall, preconditioning enhanced expression of the established potency marker *IDO* and produced condition-dependent changes in the candidate biomarkers *CTSS* and *C1s* following inflammatory challenge. These responses were generally more consistent in UC MSCs and were detectable as early as 6 h after thaw.

## DISCUSSION

Licensing MSCs to potentiate their therapeutic benefits^11,24–26,31,43^ and to limit variability^49,50^ is an increasingly popular approach. While it has been reported that MSCs with high glycolytic flux are functionally superior for therapeutic use^16,40^, few studies have examined how cytokine activation modulates the metabolic status of MSCs. Our study reveals a previously undefined complexity in metabolic preconditioning strategies for MSC therapies, whereby conditions conventionally selected to enhance immunomodulatory activity may not be optimal for metabolic activation. Activation with IFN-γ or TNF-α for 2 hours was sufficient to shift MSCs toward glycolysis, whereas responses after 24 hours were more heterogeneous and were accompanied by reduced respiratory activity in several donor populations. Since IFN-γ and TNF-α licensing protocols typically dose cells for 24 to 72 hours^11,24,27,31,35,43^, our results indicate that prolonged licensing does not necessarily confer a corresponding metabolic advantage and that metabolic state should be considered independently when optimizing preconditioning protocols. Many studies have shown that combining IFN-γ and TNF-α has a synergistic effect on the immunomodulatory response of MSCs. We found that combined IFN-γ and TNF-α stimulation produced a different metabolic response from stimulation with either cytokine alone, increasing glycolytic flux while also increasing maximal respiration and spare respiratory capacity in most donor populations. Thus, cytokine activation did not simply switch MSCs from oxidative phosphorylation to glycolysis, but could increase metabolic capacity across both pathways. Notably, these responses were detectable after short exposure and at cytokine concentrations substantially below those commonly used for MSC licensing. This differs from conventional licensing strategies, which have largely been optimized around induction of immunomodulatory mediators rather than cellular energetics. Our results therefore suggest that the cytokine concentration and exposure time required to alter MSC metabolic state may differ from those required to maximize expression of individual immunomodulatory factors. Defining the minimum cytokine exposure required to establish a therapeutically favorable metabolic and functional phenotype could reduce manufacturing time and cytokine use while limiting unnecessary prolonged stimulation.

While the majority of MSC donor populations behaved similarly, some exceptions were documented. For example, UC2 had reduced spare respiratory capacity after only 2 hours of stimulation with either IFN-γ or TNF-α, an effect that was not seen in the other populations. After 24 hours of TNF-α exposure, UC2 exhibited increased basal respiration and spare capacity, with a corresponding rise in ATP-linked respiration, all of which were reduced in the other MSC populations in the same conditions. Interestingly, UC2 was one of the strongest responders to cytokine challenge with respect to expression of *IDO*, *C1s* and *CTSS*. Whether UC2 has different functional potency compared to the other donor populations is not yet known. Nevertheless, these donor-specific responses reinforce previous observations that MSC populations differ substantially in their responsiveness to inflammatory licensing and that increasing cytokine exposure does not necessarily overcome weak donor responsiveness^46^. Thus, a single licensing protocol may not produce a uniform metabolic or functional state across MSC products.

Many important, open questions remain about how MSC performance *in vitro* correlates with their performance *in vivo*, particularly with respect to thresholding predictive biomarkers^8,51,52^. For example, levels of *IDO* expression by IFN-γ-licensed MSCs have been linked with their immunosuppressive potency^53^. However, Boyt et al. showed that *IDO* expression and kynurenine production could be elevated by increasing IFN-γ concentrations and the duration of exposure, but that these biochemical increases did not reliably correlate with enhanced functional outcomes (e.g., PBMC suppression)^46^. More broadly, surrogate potency markers must ultimately be linked to a relevant biological activity, and no single measurement is expected to capture the multiple mechanisms contributing to MSC therapeutic function. Our observation that pronounced metabolic responses did not consistently coincide with stronger potency-associated gene expression further supports this distinction. Rather than identifying glycolytic flux as a direct surrogate for immunomodulatory potency, our findings suggest that metabolic fitness and cytokine responsiveness represent related but non-equivalent attributes of MSC quality. Boyt et al. also reported that some MSC donors responded robustly to licensing and exhibited strong immunosuppressive activity, while others were weak responders; these differences could not be overcome by increasing IFN-γ exposure^46^. This agrees with our own data, which revealed donor-linked variability in the metabolic responses to various doses and durations of cytokine stimulation. These findings underscore the significant challenge of MSC heterogeneity and suggest that a “one-size-fits-all” preconditioning approach may not be feasible for manufacturing MSC products.

Numerous reports have highlighted a transient functional impairment of cryopreserved MSCs after thaw, necessitating a “refresh” step that allows the cells to recover in culture before administration^11,13,17,54^. Consistent with this phenomenon, in this study all MSC populations exhibited signs of metabolic impairment after thaw. Licensing has been shown to improve the functional potency of thawed MSCs, so we hypothesized that metabolic function might similarly be improved. For example, IFN-γ pre-licensing before cryopreservation has been shown to restore post-thaw suppression of T-cell proliferation and cytotoxic T-cell degranulation, although it did not fully rescue cryopreservation-associated defects in MSC lung tropism^11^. Thus, preconditioning may preserve selected post-thaw functions without universally restoring MSC fitness.

Mitochondrial respiration is known to be impaired post-thaw, which can result in a compensatory upregulation of glycolysis^7,11,55^. This compensatory mechanism is evident in the unconditioned, thawed MSCs like UC2, which had the highest ECAR value combined with the lowest mitochondrial respiration parameters. Preconditioning UC2 specifically enhanced aerobic respiration without elevating ECAR compared to fresh or unconditioned counterparts, suggesting that preconditioning restored mitochondrial metabolic capacity without a corresponding increase in glycolytic flux. Similar improvement was only documented for two other donor populations – BM1 and BM2. Taken together, these data suggest that preconditioning may facilitate post-thaw metabolic recovery, but that this benefit is not universal across MSC populations. Further work is necessary to identify MSC attributes that make them amenable to metabolic preconditioning and to correlate metabolic status with therapeutic performance.

Importantly, preconditioning also influenced how rapidly thawed MSCs responded to a subsequent inflammatory challenge. In UC MSCs, preconditioned cells showed enhanced *IDO* expression after short post-thaw recovery, indicating that preconditioning can preserve or accelerate an established molecular response to inflammatory stimulation. The candidate biomarkers *CTSS* and *C1s* also detected these early preconditioning-associated responses in both UC donors, supporting their potential utility as complementary measures of MSC responsiveness. Their performance was less consistent in BM MSCs: *CTSS* detected selected preconditioning-associated responses, whereas *C1s* showed comparatively little discrimination between preconditioned and unconditioned thawed BM cells. Thus, these data provide initial evidence that *CTSS* and *C1s* can report changes in post-thaw cytokine responsiveness, while also demonstrating that their utility may depend on MSC tissue source, donor, and assay conditions. Further characterization against established functional assays will be necessary before either marker can be considered a surrogate of MSC potency.

We also explored the hypothesis that metabolically superior (i.e. more glycolytic) MSCs may also be more functionally responsive, but did not observe a clear association. For example, all 6 metabolic parameters of preconditioned BM2 were enhanced by 6 hours after thaw. Despite this strong metabolic response to preconditioning, expression of the potency-associated genes *IDO* and *CTSS* was not uniformly improved following cytokine challenge. In contrast, UC1 was the only population for which preconditioning provided limited metabolic benefit after thaw, yet it exhibited the highest expression levels of the potency biomarkers 6 hours after thaw. These discordant responses argue against using metabolic flux alone as a proxy for immunomodulatory potency. Instead, metabolic fitness and inducible molecular responses may capture complementary dimensions of MSC quality. A multidimensional assessment incorporating cellular energetics together with rapidly inducible molecular biomarkers such as *IDO* and, pending further validation, *CTSS* and *C1s*, may therefore provide a more informative framework for evaluating MSC fitness following manufacturing and cryopreservation. Such an approach is consistent with emerging potency-assay strategies that combine complementary measures of MSC quality and function rather than relying on a single surrogate endpoint^56,57^.

Collectively, these findings demonstrate that cytokine dose, duration, and combination can differentially shape MSC metabolic state, and that preconditioning can improve both metabolic recovery and early responsiveness to inflammatory challenge after thaw in a donor-dependent manner. Importantly, metabolic recovery did not consistently parallel potency-associated gene expression, indicating that these measures capture complementary rather than interchangeable aspects of MSC fitness. The ability of the candidate biomarkers *CTSS* and *C1s* to detect preconditioning-associated responses, particularly in UC MSCs, further supports their continued evaluation alongside established markers such as *IDO*. Together, these findings provide a framework for refining MSC preconditioning strategies and developing multidimensional approaches to assess cell fitness following manufacturing and cryopreservation.

## Data Availability

The authors confirm that the data supporting the findings of this study are available within the article and its supplementary materials.

## Acknowledgements

We thank Tissue Regeneration Therapeutics Inc. (TRT), Toronto, Canada, for provision of human umbilical cord perivascular cells. The initial studies underlying this work were conducted at Aurora BioSolutions Inc. The study was subsequently expanded and the principal experimental work completed in the laboratory of L.R.B. at Simon Fraser University. H.B. was supported by a Stem Cell Network Summer Studentship, an NSERC Undergraduate Student Research Award and a CIHR Canada Graduate Scholarship – Master’s award. This work was funded by a Natural Sciences and Engineering Research Council of Canada Discovery Grant and a Canada Research Chair in MSC Biology awarded to L.R.B.

## Author Contributions

L.R.B. conceived the original study and supervised its subsequent development. B.T.D. performed the initial studies that established the experimental basis for this work and generated the data reported in Figure S1. H.B. contributed substantially to the experimental design and methodology, performed the experiments reported in the main manuscript, and analyzed the data. L.R.B. and H.B. interpreted the results and prepared the manuscript. B.T.D. contributed to data interpretation and revision of the manuscript. All authors reviewed and approved the final manuscript.

## References

1. Balci, D. & Can, A. The Assessment of Cryopreservation Conditions for Human Umbilical Cord Stroma-Derived Mesenchymal Stem Cells towards a Potential Use for Stem Cell Banking. Curr Stem Cell Res Ther 8, 60–72 (2013).

2. Biobanking and Cryopreservation of Stem Cells. Vol 951. (Springer International Publishing, 2016). doi:10.1007/978-3-319-45457-3.

3. Cottle, C., Porter, A. P. & Lipat, A. Impact of Cryopreservation and Freeze-Thawing on Therapeutic Properties of Mesenchymal Stromal/Stem Cells and Other Common Cellular Therapeutics. Curr Stem Cell Rep 8, 72–92 (2022).

4. Chabot, D., Tremblay, T., Paré, I., Bazin, R. & Loubaki, L. Transient warming events occurring after freezing impairs umbilical cord–derived mesenchymal stromal cells functionality. Cytotherapy 19, 978–989 (2017).

5. Maughon, T. S., Shen, X. & Huang, D. Metabolomics and cytokine profiling of mesenchymal stromal cells identify markers predictive of T-cell suppression. Cytotherapy 24, 137–148 (2022).

6. Davies, O. G., Smith, A. J., Cooper, P. R., Shelton, R. M. & Scheven, B. A. The effects of cryopreservation on cells isolated from adipose, bone marrow and dental pulp tissues. Cryobiology 69, 342–347 (2014).

7. Galipeau, J. Concerns arising from MSC retrieval from cryostorage and effect on immune suppressive function and pharmaceutical usage in clinical trials. ISBT Sci Ser 8, 100–101 (2013).

8. Lu, W. & Allickson, J. Mesenchymal stromal cell therapy: Progress to date and future outlook. Mol. Ther. J. Am. Soc. Gene Ther. 33, 2679–2688 (2025).

9. Prockop, D. J. & Youn Oh, J. Mesenchymal Stem/Stromal Cells (MSCs): Role as Guardians of Inflammation. Mol. Ther. 20, 14–20 (2012).

10. Nystedt, J. The Utilization of Freezing Steps in Mesenchymal Stromal Cell (MSC) Manufacturing: Potential Impact on Quality and Cell Functionality Attributes. Front Immunol 10, (2019).

11. Chinnadurai, R. et al. Cryopreserved Mesenchymal Stromal Cells Are Susceptible to T-Cell Mediated Apoptosis Which Is Partly Rescued by IFNγ Licensing. Stem Cells Dayt. Ohio 34, 2429–2442 (2016).

12. Pollock, K., Sumstad, D., Kadidlo, D., McKenna, D. H. & Hubel, A. Clinical mesenchymal stromal cell products undergo functional changes in response to freezing. Cytotherapy 17, 38–45 (2015).

13. François, M. et al. Cryopreserved mesenchymal stromal cells display impaired immunosuppressive properties as a result of heat-shock response and impaired interferon-γ licensing. Cytotherapy 14, 147–152 (2012).

14. Haack-Sørensen, M. & Kastrup, J. Cryopreservation and Revival of Mesenchymal Stromal Cells. Methods Mol. Biol. Humana Press 2011, 161–174.

15. Moll, G., Alm, J. J. & Davies, L. C. Do Cryopreserved Mesenchymal Stromal Cells Display Impaired Immunomodulatory and Therapeutic Properties? Stem Cells 32, 2430–2442 (2014).

16. Yuan, X., Logan, T. M. & Ma, T. Metabolism in Human Mesenchymal Stromal Cells: A Missing Link Between hMSC Biomanufacturing and Therapy? Front Immunol 10, (2019).

17. Bahsoun, S., Coopman, K. & Akam, E. C. The impact of cryopreservation on bone marrow-derived mesenchymal stem cells: a systematic review. J Transl Med 17, (2019).

18. Gramlich, O. W. et al. Cryopreserved Mesenchymal Stromal Cells Maintain Potency in a Retinal Ischemia/Reperfusion Injury Model: Toward an off-the-shelf Therapy. Sci Rep 6, (2016).

19. Cruz, F. F., Borg, Z. D. & Goodwin, M. Freshly Thawed and Continuously Cultured Human Bone Marrow-Derived Mesenchymal Stromal Cells Comparably Ameliorate Allergic Airways Inflammation in Immunocompetent Mice. Stem Cells Transl Med 4, 615–624 (2015).

20. Hoogduijn, M. J., Witte, S. F. H. & Luk, F. Effects of Freeze–Thawing and Intravenous Infusion on Mesenchymal Stromal Cell Gene Expression. Stem Cells Dev 25, 586–597 (2016).

21. Yan, B., Chen, H., Yan, L., Yuan, Q. & Guo, L. Cryopreserved Umbilical Cord Mesenchymal Stem Cells Show Comparable Effects to Un-Cryopreserved Cells in Treating Osteoarthritis. Cell Transplant. 34, 09636897241297631 (2025).

22. Linkova, D. D., Rubtsova, Y. P. & Egorikhina, M. N. Cryostorage of Mesenchymal Stem Cells and Biomedical Cell-Based Products. Cells 11, 2691 (2022).

23. Song, Y. et al. Optimizing therapeutic outcomes: preconditioning strategies for MSC-derived extracellular vesicles. Front. Pharmacol. 16, 1509418 (2025).

24. Li, H., Ji, X.-Q., Zhang, S.-M. & Bi, R.-H. Hypoxia and inflammatory factor preconditioning enhances the immunosuppressive properties of human umbilical cord mesenchymal stem cells. World J. Stem Cells 15, 999–1016 (2023).

25. Tunstead, C. et al. The ARDS microenvironment enhances MSC-induced repair via VEGF in experimental acute lung inflammation. Mol. Ther. 32, 3422–3432 (2024).

26. Hawthorne, I. J. et al. Human macrophage migration inhibitory factor potentiates mesenchymal stromal cell efficacy in a clinically relevant model of allergic asthma. Mol. Ther. 31, 3243–3258 (2023).

27. Soltero-Rivera, M., Arzi, B., Bourebaba, L. & Marycz, K. Impact of Pro-Inflammatory Cytokine Preconditioning on Metabolism and Extracellular Vesicles in Feline Mesenchymal Stromal Cells: A Preliminary Study. Stem Cells Publ. Online April 1, (2025).

28. Caffi, V. et al. Pre-conditioning of Equine Bone Marrow-Derived Mesenchymal Stromal Cells Increases Their Immunomodulatory Capacity. Front. Vet. Sci. 7, 318 (2020).

29. Mendt, M., Daher, M. & Basar, R. Metabolic Reprogramming of GMP Grade Cord Tissue Derived Mesenchymal Stem Cells Enhances Their Suppressive Potential in GVHD. Front Immunol 12, (2021).

30. Xu, C. et al. TNFα and IFNγ rapidly activate PI3K-AKT signaling to drive glycolysis that confers mesenchymal stem cells enhanced anti-inflammatory property. Stem Cell Res. Ther. 13, 491 (2022).

31. Xu, C. et al. TNFα and IFNγ rapidly activate PI3K-AKT signaling to drive glycolysis that confers mesenchymal stem cells enhanced anti-inflammatory property. Stem Cell Res. Ther. 13, 491 (2022).

32. Liu, Y., Yuan, X., Muñoz, N., Logan, T. M. & Ma, T. Commitment to Aerobic Glycolysis Sustains Immunosuppression of Human Mesenchymal Stem Cells. Stem Cells Transl Med 8, 93–106 (2019).

33. Jacques, V. et al. Metabolic conditioning enhances human bmMSC therapy of doxorubicin-induced heart failure. Stem Cells 42, 874–888 (2024).

34. Li, H., Dai, H. & Li, J. Immunomodulatory properties of mesenchymal stromal/stem cells: The link with metabolism. J. Adv. Res. 45, 15–29 (2023).

35. Yao, M. et al. Cross talk between glucose metabolism and immunosuppression in IFN-γ–primed mesenchymal stem cells. Life Sci. Alliance 5, e202201493 (2022).

36. Tonarova, P., Lochovska, K., Pytlik, R. & Hubalek Kalbacova, M. The Impact of Various Culture Conditions on Human Mesenchymal Stromal Cells Metabolism. Stem Cells Int. 2021, 1–15 (2021).

37. Lavrentieva, A., Majore, I., Kasper, C. & Hass, R. Effects of hypoxic culture conditions on umbilical cord-derived human mesenchymal stem cells. Cell Commun. Signal. 8, 18 (2010).

38. Estrada, J. C. et al. Culture of human mesenchymal stem cells at low oxygen tension improves growth and genetic stability by activating glycolysis. Cell Death Differ. 19, 743–755 (2012).

39. Buravkova, L. B., Andreeva, E. R., Gogvadze, V. & Zhivotovsky, B. Mesenchymal stem cells and hypoxia: Where are we? Mitochondrion 19, 105–112 (2014).

40. Contreras-Lopez, R. et al. The ATP synthase inhibition induces an AMPK-dependent glycolytic switch of mesenchymal stem cells that enhances their immunotherapeutic potential. Theranostics 11, 445–460 (2021).

41. Horie, S. et al. Cytokine pre-activation of cryopreserved xenogeneic-free human mesenchymal stromal cells enhances resolution and repair following ventilator-induced lung injury potentially via a KGF-dependent mechanism. Intensive Care Med. Exp. 8, 8 (2020).

42. Rossello-Gelabert, M., Igartua, M., Santos-Vizcaino, E. & Hernandez, R. M. Fine-tuning licensing strategies to boost MSC-based immunomodulatory secretome. Stem Cell Res. Ther. 16, 183 (2025).

43. López-García, L. & Castro-Manrreza, M. E. TNF-α and IFN-γ Participate in Improving the Immunoregulatory Capacity of Mesenchymal Stem/Stromal Cells: Importance of Cell-Cell Contact and Extracellular Vesicles. Int. J. Mol. Sci. 22, 9531 (2021).

44. Li, W., Liu, Q., Shi, J., Xu, X. & Xu, J. The role of TNF-α in the fate regulation and functional reprogramming of mesenchymal stem cells in an inflammatory microenvironment. Front. Immunol. 14, 1074863 (2023).

45. Egea, V. et al. TNF-α respecifies human mesenchymal stem cells to a neural fate and promotes migration toward experimental glioma. Cell Death Differ. 18, 853–863 (2011).

46. Boyt, D. T., Boland, L. K., Burand, A. J., Brown, A. J. & Ankrum, J. A. Dose and duration of interferon γ pre-licensing interact with donor characteristics to influence the expression and function of indoleamine-2,3-dioxygenase in mesenchymal stromal cells. J. R. Soc. Interface 17, 20190815 (2020).

47. Wang, J. et al. Immunomodulatory potential of cytokine-licensed human bone marrow-derived mesenchymal stromal cells correlates with potency marker expression profile. Stem Cells 42, 1040–1054 (2024).

48. Wang, J. et al. Immunomodulatory potential of cytokine-licensed human bone marrow-derived mesenchymal stromal cells correlates with potency marker expression profile. Stem Cells 42, 1040–1054 (2024).

49. Rossello-Gelabert, M., Igartua, M., Santos-Vizcaino, E. & Hernandez, R. M. Fine-tuning licensing strategies to boost MSC-based immunomodulatory secretome. Stem Cell Res. Ther. 16, 183 (2025).

50. Wiese, D. M., Wood, C. A., Ford, B. N. & Braid, L. R. Cytokine Activation Reveals Tissue-Imprinted Gene Profiles of Mesenchymal Stromal Cells. Front. Immunol. 13, 917790 (2022).

51. Viswanathan, S. & Galipeau, J. Hallmarks of MSCs: Key quality attributes for pharmacology and clinical use. Cell Stem Cell 32, 878–894 (2025).

52. Renesme, L. et al. Delphi-driven consensus definition for mesenchymal stromal cells and clinical reporting guidelines for mesenchymal stromal cell-based therapeutics. Cytotherapy 27, 146–168 (2025).

53. François, M., Romieu-Mourez, R., Li, M. & Galipeau, J. Human MSC Suppression Correlates With Cytokine Induction of Indoleamine 2,3-Dioxygenase and Bystander M2 Macrophage Differentiation. Mol. Ther. 20, 187–195 (2012).

54. Braid, L. R., Wood, C. A., Wiese, D. M. & Ford, B. N. Intramuscular administration potentiates extended dwell time of mesenchymal stromal cells compared to other routes. Cytotherapy 20, 232–244 (2018).

55. Abazari, A., Hawkins, B. J., Clarke, D. M. & Mathew, A. J. Biopreservation Best Practices: A Cornerstone in the Supply Chain of Cell-based Therapies – MSC Model Case Study. Cell Gene Ther. Insights 3, 853–871 (2017).

56. Niekamp, P., Gu, D., Jiang, J., Woods, E. J. & Johnstone, B. H. Development and validation of a potency assay matrix for optimized and consistent manufacture of clinical mesenchymal stem/stromal cells. Front. Immunol. 17, 1725191 (2026).

57. Galipeau, J. et al. International Society for Cellular Therapy perspective on immune functional assays for mesenchymal stromal cells as potency release criterion for advanced phase clinical trials. Cytotherapy 18, 151–159 (2016).

